# DIV-1/PolA2 Promotes GLP-1/Notch-Mediated Cellular Events in *Caenorhabditis elegans*

**DOI:** 10.1101/088708

**Authors:** Dong Suk Yoon, Dong Seok Cha, Myon-Hee Lee

## Abstract

Notch signaling is a highly conserved cell signaling system in most multicellular organisms and plays a critical role in animal development. In various tumor cells, Notch signaling is elevated and has been considered as an important target in cancer treatments. In *C. elegans*, GLP-1 (one of two *C. elegans* Notch receptors) activity is required for cell fate specification in germline and somatic tissues. In this study, we have identified *div-1* gene as a positive regulator for GLP-1/Notch-mediated cellular events. *C. elegans div-1* encodes the B subunit of the DNA polymerase alpha-primase complex and is highly expressed in proliferative germ cells. Functional analyses demonstrated that *i*) DIV-1 is required for the robust proliferation typical of the germline, *ii*) loss of DIV-1 enhances and suppresses specific phenotypes that are associated with reduced and elevated GLP-1/Notch activity in germline and somatic tissues, and *iii*) DIV-1 works together with FBF/PUF proteins, downstream regulators of GLP-1/Notch signaling, to promote germline stem cell (GSC) maintenance and germline proliferation. To maintain GSCs and proliferative cell fate, GLP-1/Notch activity must remain above a threshold for proliferation/differentiation decision. Our results propose that DIV-1 may control the level of threshold for GLP-1/Notch-mediated germline proliferation. PolA2, a mammalian homolog of the *C. elegans* DIV-1, has been emerged as a therapeutic target for non-small cell lung cancer (NSCLC). Notably, Notch signaling is altered in approximately one third of NSCLCs. Therefore, the discovery of the DIV-1 effect on GLP-1/Notch-mediated cellular events has implications for our understanding of vertebrate PolA2 protein and its influence on stem cell maintenance and tumorigenesis.

## INTRODUCTION

Germline stem cells (GSCs) are characterized by their ability to produce themselves (known as “self-renewal”) and to generate gametes – sperm or eggs (known as “differentiation”). A balance between self-renewal and differentiation of GSCs is strictly regulated by a systematic regulatory network, including extrinsic cues and intrinsic regulators (Wong et al., 2005). Therefore, aberrant regulation of this network can result in either loss of a specific germ cell type or over-proliferation of undifferentiated germ cells, which is associated with germline tumors (Wong et al., 2005). One of the key extrinsic cues is Notch signaling (Liu et al., 2010). This signaling plays varied and essential roles in regulating many types of stem cells (Liu et al., 2010). In *C. elegans* germline, GLP-1 (one of two *C. elegans* Notch receptors) signaling promotes mitotic proliferation of germ cells and maintenance of GSCs (Kimble and Crittenden, 2007) (Fig. 1A). Briefly, Notch ligand, LAG-2, is expressed in stem cell niche (called by “distal tip cell [DTC]” in *C. elegans*) (Henderson et al., 1994) and interacts with the GLP-1/Notch receptor, following by proteolytic cleavage of the GLP-1/Notch receptor. GLP-1/Notch intracellular domain (NICD) is translocated from membrane into nucleus. In nucleus, the NICD forms ternary complex with LAG-1/CSL DNA binding protein and LAG-3/SEL-8/Mastermind transcription coactivator to activate the expression of target genes (Fig. 1B and 1C). Those include *fbf-2* (a member of the PUF (Pumilio/FBF) RNA-binding protein family) (Lamont et al., 2004), *lip-1* (a homolog of the dual-specificity phosphatase) (Lee et al., 2006), *lst-1* (lateral signaling target-1, unknown protein) (Kershner et al., 2014), and *sygl-1* (synthetic Glp-1, unknown protein) (Kershner et al., 2014). These genes function redundantly to maintain GSCs in *C. elegans* germline (Kershner et al., 2014). For example, *lst-1* and *sygl-1* single mutants possessed germlines comparable in size and organization to wild-type, and they were self-fertile (Kershner et al., 2014). However, most *lst-1 sygl-1* double mutants displayed a premature meiotic entry (called Glp [GermLine proliferation defect], no mitotic cells, few germ cells, and all sperm) phenotype and they were thus sterile (Kershner et al., 2014). In addition, FBF-2 and LIP-1 proteins promote germline proliferation by inhibiting the expression or activity of the meiosis-promoting regulators (e.g., GLD-1/Quaking and MPK-1/ERK) (Lee et al., 2006; Kimble and Crittenden, 2007) and cell cycle regulators (e.g., CKI-2, a Cyclin E/CDK2 inhibitor (Kalchhauser et al., 2011)) in the *C. elegans* germline. Notably, Cyclin E has been identified as a positive regulator for GSC maintenance and germline proliferation in *C. elegans* (Fox et al., 2011) and *Drosophila* (Ables and Drummond-Barbosa, 2013). Therefore, elevated GLP-1/Notch activity promotes germline proliferation and inhibits meiotic entry, resulting in germline tumors (Berry et al., 1997). However, reduced GLP-1/Notch activity causes all GSCs to cease self-renewal/germline proliferation and differentiate as sperm (Glp phenotype) (Fig. 1D and 1E).

**Figure 1.**
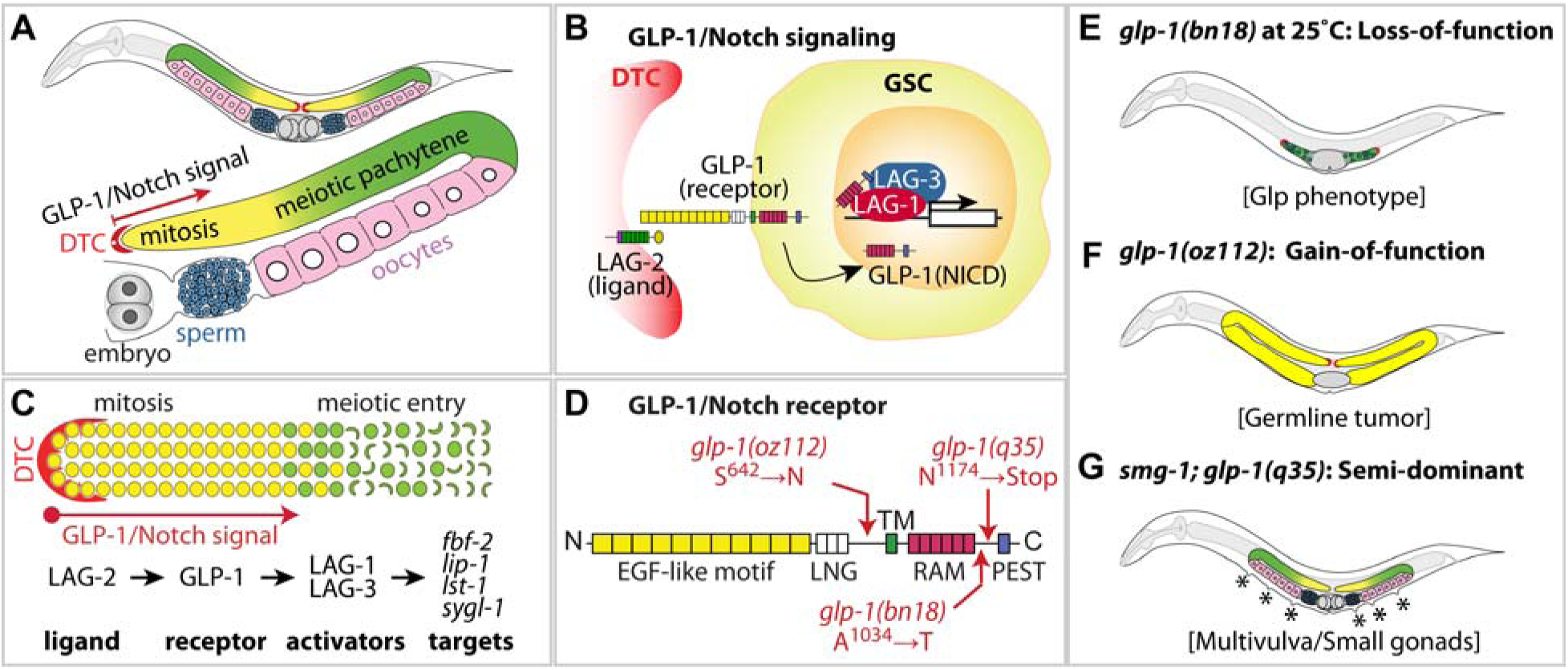
*C. elegans* germline and GLP-1/Notch signaling pathway. (A) Schematic of adult hermaphrodite gonad. Germ cells at the distal end of the germline, including germline stem cells (GSCs), divide mitotically (yellow). As germ cells move proximally, they enter meiosis (green) and differentiate into either sperm (blue) or oocytes (pink). GLP-1/Notch signaling is activated in the distal mitotic germline. Slightly modified from (Kobet et al., 2014). (B) *C. elegans* GLP-1/Notch signaling pathway. The LAG-2 ligand, localized to the DTC, signals to GLP-1/Notch receptor in GSCs and mitotically dividing germ cells. Upon GLP-1 activation, the GLP-1 intracellular domain (ICD), LAG-1 and LAG-3 form a ternary complex in the nucleus and activate transcription of target genes. Slightly modified from (Kobet et al., 2014). (C) GLP-1/Notch signaling pathway in the *C. elegans* distal germline. Red, DTC; yellow circles, germ cells in mitotic cell cycle; green circles, germ cells in meiotic S phase; green crescents, germ cells in early meiotic prophase. Bottom shows the core regulators of GLP-1/Notch signaling pathway and its target genes. (D) The GLP-1 domains and mutations. *glp-1(bn18)* is a temperature-sensitive, loss-of-function mutation (A1034➔T) (Kodoyianni et al., 1992), *glp-1(oz112)* is a gain-of-function mutation (S642➔N) (Berry et al., 1997), and *glp-1(q35)* is a nonsense allele with both gain-of-function in somatic vulva precursor cells and loss-of-function in germline (Mango et al., 1991). EGF, epidermal growth factor; LNG, LIN-12/Notch/GLP-1; TM, transmembrane; RAM, Rbp-associated molecule domain; PEST, Proline (P) Glutamic acid (E), Serine (S), and Threonine (T). (E-G) Schematics of germline phenotypes in adult hermaphrodites that are associated with *glp-1(bn18)* loss-of-function (E), *glp-1(oz112)* gain-of-function (F), and *smg-1; glp-1(q35)* semi-dominant mutations (G).

Cell cycle control is a critical step to generate specific cells and tissues during developments and maintain cellular homeostasis in adult (Besson et al., 2008). The cell cycle is tightly regulated by complexes containing Cyclins and Cyclin-dependent kinases (CDKs) (Johnson and Walker, 1999; Murray, 2004). Dysregulation of this process can result in either loss of specific cell types or over-proliferation, which could lead to a number of human diseases, including cancers (Vermeulen et al., 2003). In general, cell cycle process can be divided in two stages: interphase (accumulating nutrients and duplicating DNA) and mitosis (dividing process). The interphase includes G1, S, and G2 phases. During the G1 phase, cells prepare for the process of DNA replication and make a decision to enter S phase. Cyclin E associates with CDK2, and pushes the cell from G1 to S phase (Ohtsubo et al., 1995). Replication of DNA occurs in S phase through interaction with many replication factors. The second gap phase, G2, is required for preparing cell division. To proceed from G2 to M phase, G2/M checkpoint needs to be passed by Cyclin A/B/CDK-1 complex (Girard et al., 1991). During mitosis, replicated chromosomes are segregated into separate nuclei followed by cytokinesis to form two daughter cells. Since cell division and development are tightly coordinated, it is plausible that cell cycle and DNA replication factors are critical for the process of development. Notably, mammalian stem cells have short G1 and greater than 70% of the stem cell population are in S phase (Savatier et al., 1996). However, the function of S phase in stem cell maintenance remains poorly understood. Moreover, key S phase regulators that are associated with stem cell maintenance have not yet been identified in any other model systems.

In this study, we explored the role of S phase factors in *C. elegans* GLP-1/Notch-mediated cellular events. We employed previously well-characterized three mutant alleles that have reduced or elevated GLP-1/Notch activities: *glp-1(bn18)*, *glp-1(ar202)*, and *glp-1(q35)* mutants (Fig. 1D-1G, Supplementary material Table S1). A focused RNAi screen using these mutants has identified *div-1* (a homolog of the human PolA2, the B subunit of the DNA polymerase alpha-primase complex) as a positive regulator of GLP-1/Notch signaling. Specifically, depletion of DIV-1 enhanced the Glp phenotype that is caused by reduced GLP-1/Notch activity. It also suppressed the formation of germline tumors and somatic multivulva that are caused by elevated GLP-1/Notch activity. Notch signaling and cell cycle regulators are highly conserved in *C. elegans*. Therefore, our findings may provide a powerful organism model system to explore the connection between Notch signaling and cell cycle progression, as well as have important implications for development and treatment of Notch signaling-associated human diseases, including cancer.

## RESULTS

### S phase arrest by chemical promotes a Glp phenotype in *glp-1(bn18)* mutants

To explore whether S phase progression is functionally linked with GLP-1/Notch-mediated germline proliferation in *C. elegans*, we utilized temperature sensitive (ts) *glp-1(bn18)* loss-of-function mutants (Fig. 1D), which provide a sensitized genetic background to determine the genetic connection with GLP-1/Notch signaling pathway (Fox et al., 2011). The *glp-1(bn18)* mutants are nearly wild-type germline proliferation albeit with reduced germ cell number (∼50% of wild-type at young adult stage) and self-fertile at permissive temperature (20°C) (Qiao et al., 1995; Maine et al., 2004). However, most *glp-1(bn18)* mutants display a Glp phenotype at restrictive temperature (25°C) (Kodoyianni et al., 1992) (Fig. 1E). First, to investigate the effect of S phase on GLP-1/Notch-mediated germline proliferation, we treated wild-type and *glp-1(bn18)* mutants from L1 larval stage with 40 mM hydroxyurea (HU, a DNA synthesis inhibitor, (Fig. 2A)) at 20°C. Three days later (adult stage), germline phenotype was determined by staining whole worms with DAPI (4’,6-diamidino-2-phenylindole). Wild-type adult worms did not display a Glp phenotype in both absence (-) and presence (+) of HU, as previously reported (Fox et al., 2011) (Fig. 2B). However, forced S phase arrest by HU treatment dramatically enhanced a Glp phenotype up to 93% in *glp-1(bn18)* mutants, while only 2% of *glp-1(bn18)* (HU-) mutants showed the Glp phenotype at the same condition (Fig. 2B). DAPI staining showed that the *glp-1(bn18)* (HU-) mutants were fertile albeit with smaller gonads (Fig. 2C), but most *glp-1(bn18)* (HU+) mutants had a typical Glp phenotype, composed solely of a few mature sperm (Fig. 2D). Notably, Fox *et al*. previously reported the treatment of *glp-1(bn18)* mutants from L4/young adult stages with HU did not induce a Glp phenotype. These results propose that S phase progression may be functionally linked with GLP-1/Notch-mediated GSC maintenance and germline proliferation during early larval stages rather than adulthood because S phase arrest by HU treatment at only early larval stages promoted the Glp phenotype.

**Figure 2.**
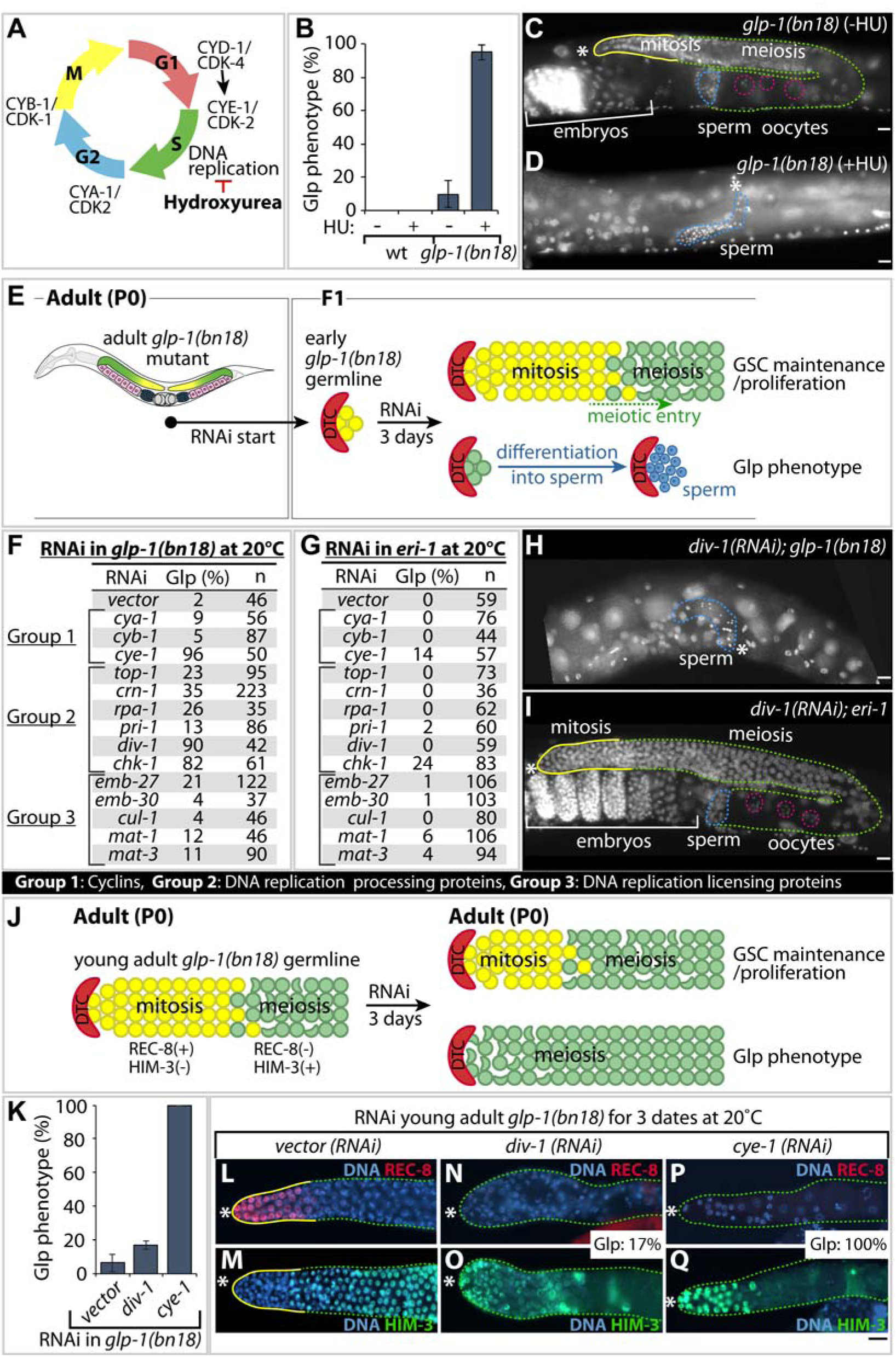
S phase progression is required for GLP-1/Notch-mediated germline proliferation. (A) Cell cycle progression and key regulators. Hydroxyurea arrests DNA replication. (B) The percentage of worms scored a Glp (premature meiotic entry) phenotype at 20°C. Standard deviation bars were calculated from three independent experiments. (C, D) DAPI-stained germlines. Solid yellow lines, mitotic germ cells; broken green lines, meiotic germ cells and differentiated gametes (broken pink circles, oocyte nuclei; broken blue line, sperm). *, distal end. (E) Schematic of RNAi experiment to test the effect of cell cycle genes on early GSC maintenance/proliferation and Glp phenotype. RNAi was performed by feeding adult staged worms (P0) bacterial expressing double stranded RNAs (dsRNAs) corresponding to the gene of interest. Phenotypes were analyzed when F1 progeny reach to adult stage (four days later). DTC, distal tip cells. (F, G) The percentage of worms scored a Glp phenotype by RNAi of cell cycle regulator genes in *glp-1(bn18)* and *eri-1(mg366)* mutants at 20°C. Group 1: RNAi of genes encoding cyclins. Group 2: RNAi of genes encoding DNA replication processing proteins. Group 3: RNAi of genes encoding DNA replication licensing proteins. The Glp phenotype was strictly defined as no mitotic cells and only sperm by DAPI staining. (H, I) DAPI-stained germlines. Solid yellow lines, mitotic germ cells; broken green lines, meiotic germ cells and differentiated gametes (broken pink circles, oocyte nuclei; broken blue line, sperm). *div-1(RNAi)* dramatically induced the Glp phenotype in *glp-1(bn18)* mutant, but not in *eri-1(mg366)* mutant. *, distal end. (J) Schematics of RNAi experiment to test the effect of *div-1*, *vector* (negative control), and *cye-1* (positive control) on adult germline proliferation. RNAi was performed by feeding young adult *glp-1 (bn18)* mutants and 3 days (72 h) later, germline phenotypes were determined by staining dissected gonads with anti-REC-8 and anti-HIM-3 antibodies. (K) The percentage of worms scored a Glp phenotype at 20°C. Standard deviation bars were calculated from three independent experiments. (L, N, P) Germline staining with anti-REC-8 antibody (proliferation marker). (M, O, Q) Germline staining with anti-HIM-3 antibody (differentiation marker). *, distal end. Scale bars: 10 µM.

### DIV-1 is required for GLP-1/Notch-mediated germline proliferation

To further explore the requirement of S phase progression in GLP-1/Notch-mediated early germline proliferation, we depleted the expression of genes, encoding Cyclins (Group 1), DNA replication processing proteins (Group 2), and DNA replication licensing proteins (Group 3) by feeding RNAi of *glp-1(bn18)* mutants (Fig. 2E-2G). Specifically, gravid adult *glp-1(bn18)* hermaphrodites were transferred to each feeding RNAi plate, and the germline phenotypes of progeny were examined by DAPI staining when they became adults (3 days after L1) (Fig. 2E). Previously, Fox *et al*., showed that GLP-1/Notch-mediated germline proliferation is required for CYE-1, but not CYA-1 and CYB-1 (Fox et al., 2011). Thus, *cye-1(RNAi)* was used as a positive control. *cya-1(RNAi)* and *cyb-1(RNAi)* were used as negative controls for focused RNAi screening (Fig. 2F). As previously reported, *cye-1(RNAi)* dramatically enhanced a Glp phenotype in *glp-1(bn18)* mutants even at 20°C, but *cya-1(RNAi)* or *cyb-1(RNAi)* did not (Fox et al., 2011) (Fig. 2F). Based on these results, we examined the effects of DNA replication regulators in Group 2 and 3 on GLP-1/Notch-mediated germline proliferation (Fig. 2F). We selected key 11 DNA replication-related genes based on Gene Ontology (GO) database (GO:0006260), previous publications, and RNAi accessibility in our library. Notably, depletion of DNA replication-related genes by RNAi enhanced the Glp phenotype in *glp-1(bn18)* mutants with a wide range of penetrance (Fig. 2F). Among them, *div-1(RNAi)* and *chk-1(RNAi)* dramatically promoted the Glp phenotype in *glp-1(bn18)* mutants even at 20°C, as seen in *cye-1(RNAi)* (Fig. 2F and 2H). We also performed RNAi experiments at the same condition in *eri-1(mg366)* mutants that is hypersensitive to RNAi (Kennedy et al., 2004) (Fig. 2G). RNAi of *cye-1* and *chk-1* genes also showed the Glp phenotype (14% and 24%, respectively) in *eri-1* mutants (Fig. 2G), but *div-1(RNAi)* did not (Fig. 2G and 2I). These results suggest that CYE-1, CHK-1, and DIV-1 play an important role in GLP-1/Notch-mediated germline proliferation. In particular, the effects of DIV-1 on germline proliferation more likely depends on GLP-1/Notch activity because *div-1(RNAi)* enhanced a Glp phenotype in *glp-1(bn18)* sensitize mutants, but not in *eri-1* mutants. Next, to ask whether *div-1* is also required for GLP-1/Notch-mediated GSC maintenance during adulthood, young adult (2.5 days after L1) staged *glp-1(bn18)* mutants were placed on the RNAi plates of *div-1*, *vector* (negative control), and *cye-1* (positive control) for 72 hours at 20°C (Fig. 2J). Their germline phenotypes were determined by staining dissected gonads with cell fate specific markers such as nucleoplasmic REC-8 for proliferative germ cells (Hansen et al., 2004b) and HIM-3 for meiotic prophase germ cells (Zetka et al., 1999) (Fig. 2L-2Q). About 5% of *vector(RNAi); glp-1(bn18)* animals displayed a Glp phenotype (Fig 2K-2M), but most *cye-1(RNAi); glp-1(bn18)* mutants had a Glp phenotype (Fig. 2K, 2N, and 2O), as previously reported (Fox et al., 2011). Interestingly, *div-1(RNAi)* at early larval stages significantly induced a Glp phenotype in *glp-1(bn18)* mutants (90%) (Fig. 2F), but *div-1(RNAi)* at adult stages slightly induced the Glp phenotype in *glp-1(bn18)* mutants (17%) (Fig. 2K, 2P, and 2Q). These results suggest that DIV-1 is required more for GLP-1/Notch-mediated early germline proliferation than for GSC maintenance during adulthood, which is consistent with the results of HU treatment in early larval stage (Fig. 2B) and adult stages (Fox et al., 2011).

### DIV-1 controls the extent of germline proliferation

To investigate the function of *div-1* in germline proliferation, we utilized a temperature sensitive (ts) *div-1(or148)* loss-of-function mutant (Encalada et al., 2000). Their germline phenotypes were analyzed by staining dissected gonads with cell fate-specific markers: anti-REC-8 (Hansen et al., 2004b) and anti-HIM-3 (Zetka et al., 1999) antibodies. The mitotic zone of the wild-type hermaphrodite gonad has two pools with distinct properties. Germ cells in the distal pool are maintained in a stem cell-like state (weak REC-8(+) signals in the nucleoplasm of mitotic nuclei, state(I)); germ cells in the proximal pool in mitotic zone are maturing from the stem cell-like state to mitotically cycling and early differentiating state (strong REC-8(+) signals on chromatin of mitotic nuclei, state(II)) (Hubbard, 2007; Fox and Schedl, 2015) (Fig. 3A). Once germ cells leave mitotic cell cycle, they enter meiotic prophase (REC-8(-)/HIM-3(+)) (Fig. 3C). Therefore, mitotic cells and meiotic cells can be distinguished using REC-8 and HIM-3 antibodies (Hubbard, 2007; Fox and Schedl, 2015). The *div-1(or148)* mutants displayed embryonic lethal phenotype due to delayed embryonic cell division at 25°C (Encalada et al., 2000). To assess the germline phenotype, we thus cultured synchronized *div-1(or148ts)* mutants until L1 at 20°C and upshift them to 25°C. Three days later (adult stage), germline phenotypes were determined by staining dissected gonads with cell fate-specific markers. Wild-type and *div-1(or148ts)* germlines grown at 25°C maintain stem cell-like state (state(I)) as well as mitotically cycling and early differentiating state (state(II)) (Fig. 3A). However, most germlines of *div-1(or148ts)* mutants grown at 25°C had no or less stem cell-like state (state(I)) (∼72%, n=42) (Fig. 3B), indicating that DIV-1 may be required for the maintenance of stem cell-like state. In addition, the germline size of *div-1(or148ts)* worms at 25°C was smaller than that of wild-type and *div-1(or148ts)* worms grown at 20°C (Fig. 3B and 3D). We then scored the number of mitotic germ cells in wild-type and *div-1(or148ts)* germlines at adult stage (three days after L1). Wild-type possess ∼225 mitotic germ cells (REC-8(+)/HIM-3(-)) (Fig. 3E). Similarly, most *div-1(or148ts)* mutants grown at 20°C are fertile and possess ∼218 mitotic germ cells (range: 200-230) (Fig. 3E). However, the average number of mitotic germ cells in the *div-1(or148ts)* mutants grown at 25°C was 109 (range: 86-140) (Fig. 3E). We next determined the S phase index (S-index) by counting number of EdU-positive nuclei in a 30-min pulse/total number of REC-8-positive nuclei, as previously reported (Crittenden et al., 2006; Fox et al., 2011). The *div-1(or148ts)* mutants grown at 25°C had longer S phase index than that grown at 20°C (60% at 20°C vs 72% at 25°C) (Fig. 3F and 3G). Moreover, adult *div-1(or148ts)* mutants grown at 25°C from L1 displayed an increased number of partially incorporated EdU(+) cells probably due to impaired DNA replication in the most distal germline (Fig. 3F and 3G). These results suggest that DIV-1 may be required for the maintenance of stem cell-like state and the normal extent of germline proliferation probably through a normal cell cycle progression.

**Figure 3.**
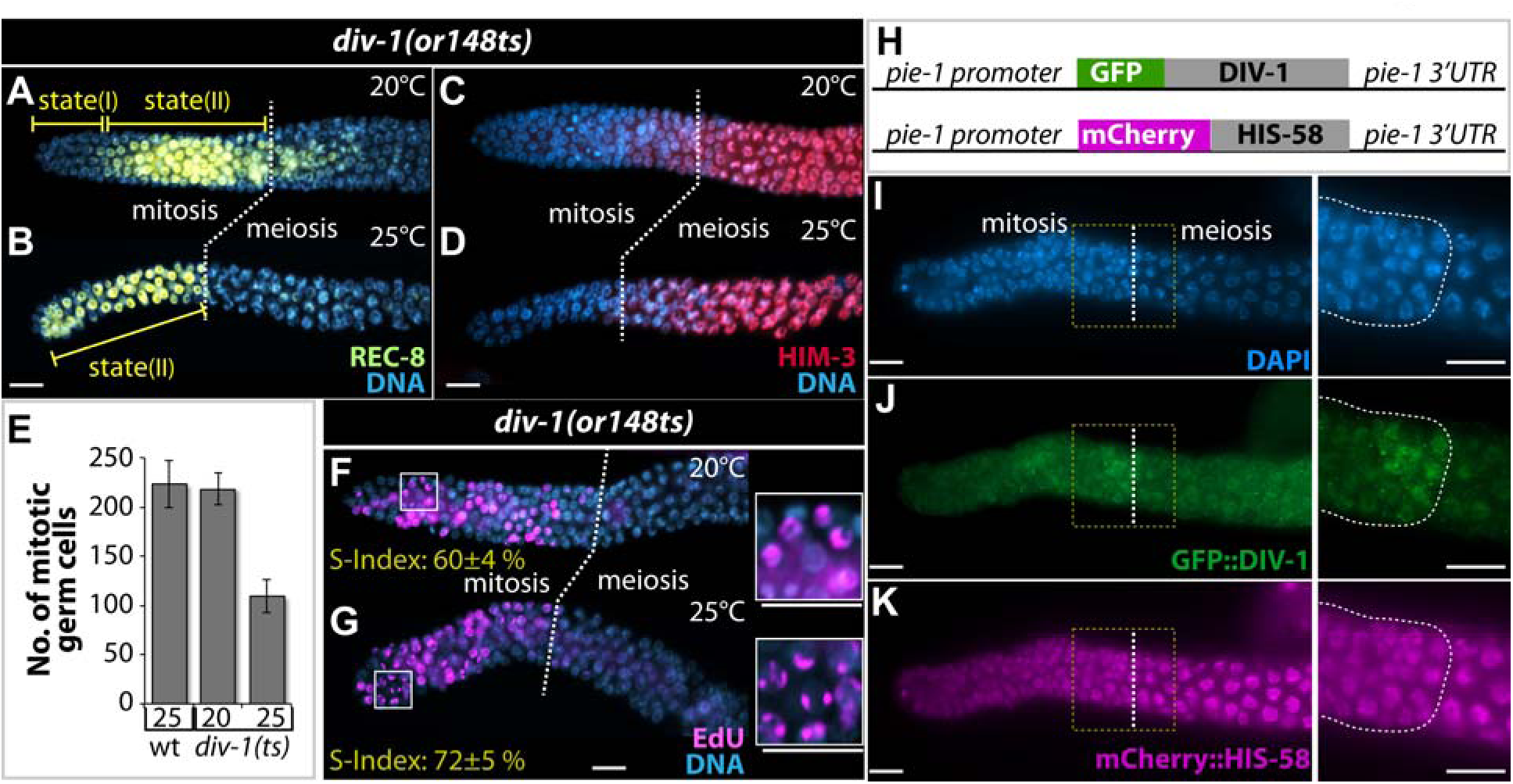
DIV-1 promotes germline proliferation. (A-D) Adult hermaphrodite germlines of *div-1(or148)* mutants grown at 20°C and 25°C were extruded and stained with REC-8 (mitotic/proliferative germ cell marker) (A and B) HIM-3 (meiotic prophase marker) (C and D). Broken lines, boundary between mitosis and meiosis. (E) Number of mitotic germ cells [REC-8(+)/HIM-3(-)]. Standard deviation bars were calculated from three independent experiments. (F, G) Adult hermaphrodite germlines of *div-1(or148)* mutants grown at 20°C and 25°C were extruded and stained with the Click-iT EdU Alexa Fluor 488 Imaging Kit (S phase maker). Broken lines, boundary between mitosis and meiosis. Magnified pictures of inset in F and G. (H) The design of the *GFP::div-1* and *mCherry::his-58* (internal control) transgenes. The *pie-1* promoter and 3’UTR are permissive for expression in all germ cells. (I-K) Expression of GFP::DIV-1 and mCherry::HIS-58 in the same germline. mCherry::HIS-58 was detected in all germ cell chromosome. GFP::DIV-1 was highly accumulated in the proximal mitotic zone containing actively dividing germ cells. Magnified pictures of inset in I-K. Broken lines, boundary between mitotic zone and meiotic prophase. Expression level of GFP::DIV-1 was dramatically decreased as germ cells enter meiotic prophase. Scale bars: 10 µM.

We next assessed the localization of DIV-1 in *C. elegans* germline. No DIV-1-specific antibody is available. Thus, the subcellular location of DIV-1 was analyzed using a transgenic worm expressing both *pie-1 (promoter)::GFP::div-1::pie-1 3’UTR* and *pie-1 (promoter)::mCherry::his-58::pie-1 3’UTR* (an internal control) transgenes (Fig. 3H). GFP::DIV-1 was detected in the nuclei of germ cells during germline development (Supplementary material Fig. S1). Notably, GFP::DIV-1 was highly accumulated in the proximal mitotic region (state(II)) that contains a mixture of mitotically cycling cells and early differentiating cells in adult germlines (Fig. 3I-3K), compared to the expression of mCherry::HIS-58). This result suggests that DIV-1 may be required for the robust proliferation typical of the mitotic germ cells during development. Next, we asked whether DIV-1 controls germline proliferation by affecting the expression of Notch signaling genes (*lag-2*, Notch ligand; *glp-1*, Notch receptor) and *gld-1* gene (a key regulator for germ cell differentiation). To answer the question, we performed *div-1(RNAi)* in wild-type worms and two transgenic worms expressing *lag-2::GFP* and *gld-1::GFP* transgenes from L1 stage and stained dissected gonads with anti-GLP-1 and anti-GFP antibodies. The staining showed that depletion of DIV-1 did not affect the expression of LAG-2, GLP-1, and GLD-1 (Supplementary material Fig. S2). This result indicates that DIV-1 may not directly influence the expression of well-known key proliferation and differentiation regulators, LAG-2, GLP-1, and GLD-1. How then does DIV-1 affect germline proliferation? One possible mechanism is that DIV-1 may promote GSC maintenance and germline proliferation by maintaining a threshold for germline proliferation/differentiation decision below GLP-1/Notch activity (see Fig. 7B and Supplementary material Fig. S5). Recent studies from Schedl’s group proposed that germline proliferation/differentiation decision is controlled by a threshold for GLP-1/Notch activity; GLP-1/Notch activity must be above a threshold for germline proliferation and its activity must be fall below a threshold for differentiation (Fox and Schedl, 2015). Based on this report, we suggest that loss of DIV-1 may rise a threshold for germline proliferation/differentiation decision (see Fig. 7B, Supplementary material Fig. S5).

### Depletion of DIV-1 partially suppresses elevated GLP-1/Notch activity-mediated germline tumor formation

Next, the role of DIV-1 in the formation of germline tumors was examined using a *glp-1(oz112)* gain-of-function (gf) mutant (see Fig. 1D and 1F). In *C. elegans*, elevated GLP-1/Notch activity promotes the formation of germline tumors (Berry et al., 1997). To explore the role of DIV-1 in the elevated GLP-1/Notch activity-mediated germline tumor formation, we depleted the expression of *div-1* gene by RNAi in *glp-1(oz112gf)* mutants (Berry et al., 1997) (Fig. 4A). While 82% of *glp-1(oz112gf)* mutants generate germline tumors at 20°C (Berry et al., 1997) (Fig. 4A and 4B), *div-1(RNAi)* partially suppressed the germline tumor formation of *glp-1(oz112gf)* mutants (∼28% reduction) (Fig. 4A and 4C). The role of DIV-1 in *glp-1(oz112gf)*-mediated germline tumor formation was confirmed in temperature-sensitive *glp-1(ar202gf)* mutants. The *glp-1(ar202gf)* mutants are typically normal at 15°C, but most of them generate proximal germline tumors at restrictive temperature (25°C) (Pepper et al., 2003). *div-1(RNAi)* also showed ∼20% and ∼10% reduction of *glp-1(ar202gf)* germline tumor formation in compared to *vector (RNAi)* at 23.5°C and 25°C, respectively (Fig. 4A). Why did the effect of *div-1(RNAi)* on *glp-1(oz112gf)* or *glp-1(ar202gf)* mutants (Fig. 4A) was much weaker than that of *div-1(RNAi)* on *glp-1(bn18)* mutants (Fig. 2E) at 20°C. One possible idea is that the phenotype of *glp-1(oz112)* or *glp-1(ar202)* mutants may be stronger than that of *glp-1(bn18)* mutants. However, we do not think because RNAi of genes that are associated with core GLP-1/Notch components (*glp-1* or *lag-3*) sufficiently repressed the formation of *glp-1(oz112gf)* germline tumors (Supplementary material Fig. S3). Moreover, RNAi of *rpa-1*, *cya-1*, and *chk-1* genes also suppressed the formation of *glp-1(ar202)* germline tumors, although RNAi of *rpa-1* and *cya-1* did not enhance a Glp phenotype in the *glp-1(bn18)* mutants (Fig. 2E, Supplementary material Fig. S3). These results also indicate that GLP-1/Notch-mediated germline proliferation and tumor formation may require different cell cycle regulators. Similarly, Nusser-Stein *et al*. recently reported that Notch signaling relies on different cell cycle regulators for the formation of a stable cell fate pattern during *C. elegans* vulva development (Nusser-Stein et al., 2012). Another possible idea is that DIV-1 is required more for germline proliferation in early larval stages than for germline proliferation during adulthood. Two lines of evidence support the latter idea: 1) the germline tumors that are caused by elevated GLP-1/Notch activity are initiated in the mid L4 stage (Pepper et al., 2003). 2) *div-1(RNAi)* from L4 staged larvae partially enhanced a Glp phenotype in adult *glp-1(bn18)* mutant germlines (Fig. 2J and 2K). Therefore, we suggest that DIV-1 is required for GLP-1/Notch-mediated GSC maintenance and germline proliferation during early larval development (Fig. 2) and also is partially necessary for the formation of germline tumors during L4 and later stages (Fig. 4).

**Figure 4.**
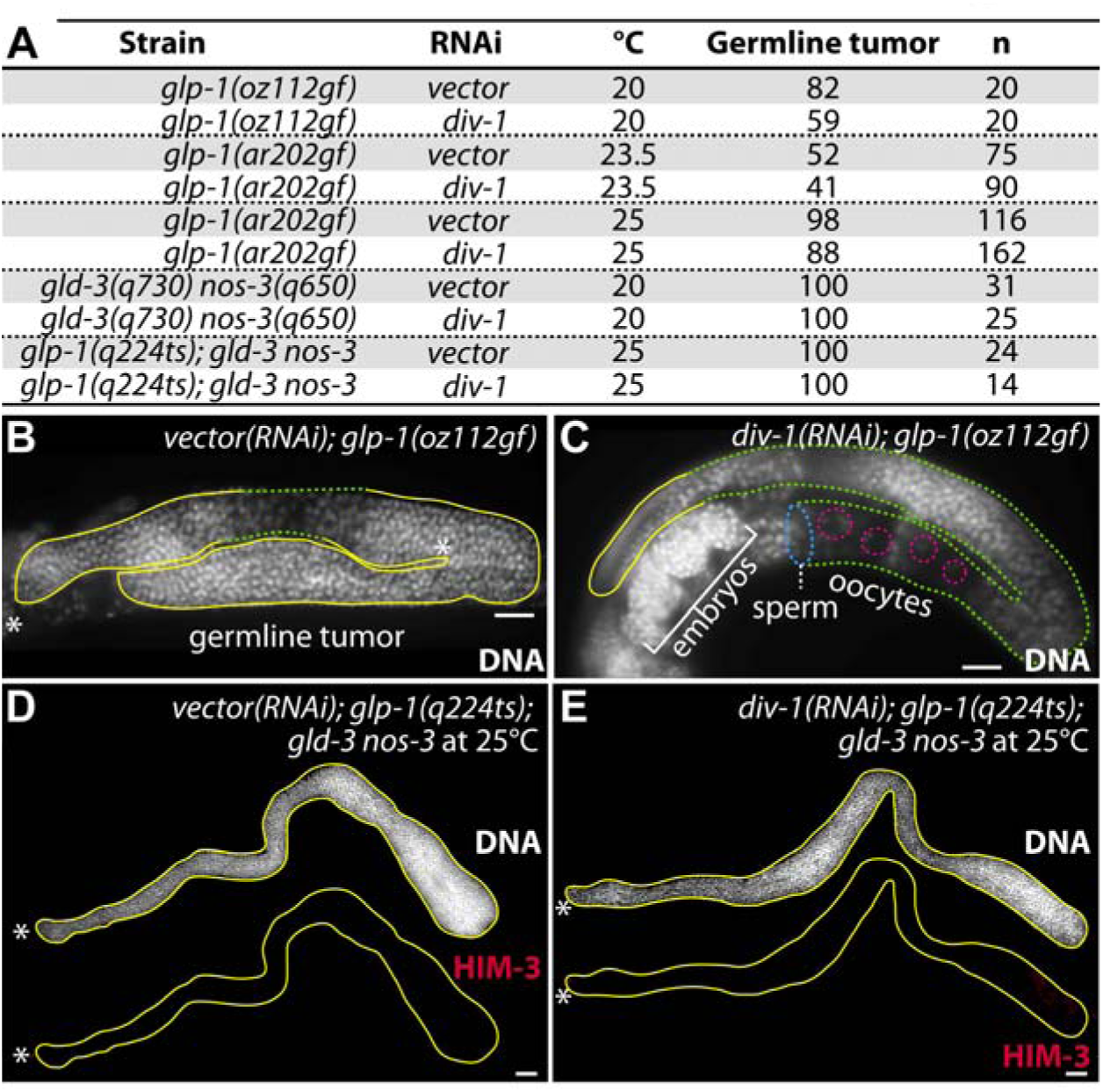
Depletion of DIV-1 partially suppresses the formation of germline tumors caused by elevated GLP-1/Notch activity. (A) The percentage of worms scored germline tumors. The germline tumors were defined as the presence of mitotically dividing cells (REC-8(+)/HIM-3(-)) in the proximal germlines. The *glp-1(oz112)* and *glp-1(ar202)* are gain-of-function mutants. The *gld-3 nos-3* mutants generate synthetic germline tumors independently with GLP-1/Notch activity. (B, C) Germline staining with DAPI. *div-1(RNAi)* partially suppressed the formation of germline tumors in *glp-1(oz112)* mutants. Solid yellow lines, mitotic germ cells; broken green lines, meiotic germ cells and differentiated gametes; (broken pink circles, oocyte nuclei; broken blue line, sperm). (D, E) Germline staining with anti-HIM-3 antibody. *div-1(RNAi)* did not promote germline differentiation in *glp-1(q224); gld-3 nos-3* mutants at 25°C. *, distal end. Scale bars: 20 µM.

Next, to ask whether the effect of DIV-1 on germline proliferation is specific to GLP-1/Notch signaling, we depleted the expression of *div-1* by RNAi in *gld-3(q730) nos-3(q650)* mutants with synthetic germline tumors. GLD-3 (a member of the Bicaudal-C family of RNA-binding proteins) and NOS-3 (a member of the Nanos family of RNA-binding proteins) promote entry into a program of differentiation (Eckmann et al., 2004). The *gld-3 nos-3* germline tumor is independent on GLP-1/Notch activity (Eckmann et al., 2004). Result showed that *div-1(RNAi)* failed to suppress *gld-3 nos-3* germline tumors (Fig. 4A). This suggests that DIV-1 may act upstream of GLD-3 and NOS-3 pathways (see Fig. 7A). To confirm this result, we also performed *div-1(RNAi)* in *glp-1(q224); gld-3 nos-3* triple mutants at 25°C. *glp-1(q224)* is a temperature-sensitive and loss-of-function mutant like *glp-1(bn18)* mutants. Most of the *glp-1(q224ts)* mutants are typically fertile, but they are completely sterile due to a Glp phenotype at 25°C (Austin and Kimble, 1987). However, homozygotes for *glp-1(q224ts); gld-3 nos-3* have germline tumors even at 25°C because the redundant GLD-3 and NOS-3 pathways act downstream of GLP-1 to promote meiotic entry (Austin and Kimble, 1987; Kadyk and Kimble, 1998; Hansen et al., 2004a) (Fig. 4A and 4D). Interestingly, *div-1(RNAi)* failed to suppress the formation of germline tumors in *glp-1(q224ts); gld-3 nos-3* mutants at 25°C (Fig. 4A and 4E). These results suggest that DIV-1 acts upstream of GLD-3 and NOS-3 pathways to GLP-1/Notch-mediated germline proliferation (see Fig. 7A).

### DIV-1 works with FBF/PUF to control mitotic cell cycle

In addition to GLP-1/Notch signaling, a battery of RNA regulators also control a balance between proliferative and differentiation. One of well conserved RNA regulators is PUF (Pumilio/FBF) RNA-binding protein. PUF proteins control various physiological processes such as stem cell maintenance and cell fate specification by interacting with 3’ untranslated regions (3’UTRs) and modulating mRNA expression in a wide variety of eukaryotes (Wickens et al., 2002). *C. elegans* has multiple PUF genes with special roles. In particular, FBF-1 and FBF-2 (96% identical, henceforth called FBF) proteins have an essential role in *C. elegans* GSC maintenance and germline proliferation (Crittenden et al., 2002). These two nearly identical FBF proteins are largely redundant: *fbf-1* and *fbf-2* single mutants are both self-fertile with germlines organized as in wild-type. By contrast, in *fbf-1 fbf-2* double mutants, GSCs are maintained until the L4 stage, but most GSCs leave mitotic cell cycle, enter meiosis, and eventually differentiated into sperm (Glp phenotype) (Crittenden et al., 2002). Interestingly, we found that the Glp phenotype of *fbf-1 fbf-2* mutants was sufficiently suppressed by additional removal of PUF-8 (another PUF protein) (Fig. 5A and 5B), a key regulator for *C. elegans* germline development (Subramaniam and Seydoux, 2003; Ariz et al., 2009; Cha et al., 2012; Racher and Hansen, 2012; Pushpa et al., 2013; Vaid et al., 2013; Datla et al., 2014; Sorokin et al., 2014; Priti and Subramaniam, 2015). The *puf-8 fbf-1 fbf-2* triple mutants displayed mitotically dividing cells in the germline (Fig. 5B). This result indicates that PUF-8 inhibits FBF-mediated GSC maintenance and germline proliferation. Intriguingly, *div-1(RNAi)* inhibited the restoration of mitotically dividing cells in *fbf-1 fbf-2 puf-8* triple mutant germlines (Fig. 5B). Mitotically dividing germ cells were detected by staining dissected gonads with mitotic/proliferative cell markers using the Click-iT EdU Alexa Fluor 488 Imaging Kit. 76% of *puf-8 fbf-1 fbf-2* germlines were positive for EdU-labeling in the germline (Fig. 5D-5G). However, only 29% of *div-1(RNAi); puf-8 fbf-1 fbf-2* germlines were positive for EdU-labeling (Fig. 5H-5K). This result suggests that DIV-1 may be required for GSC maintenance and germline proliferation in *puf-8 fbf-1 fbf-2* mutant germlines.

**Figure 5.**
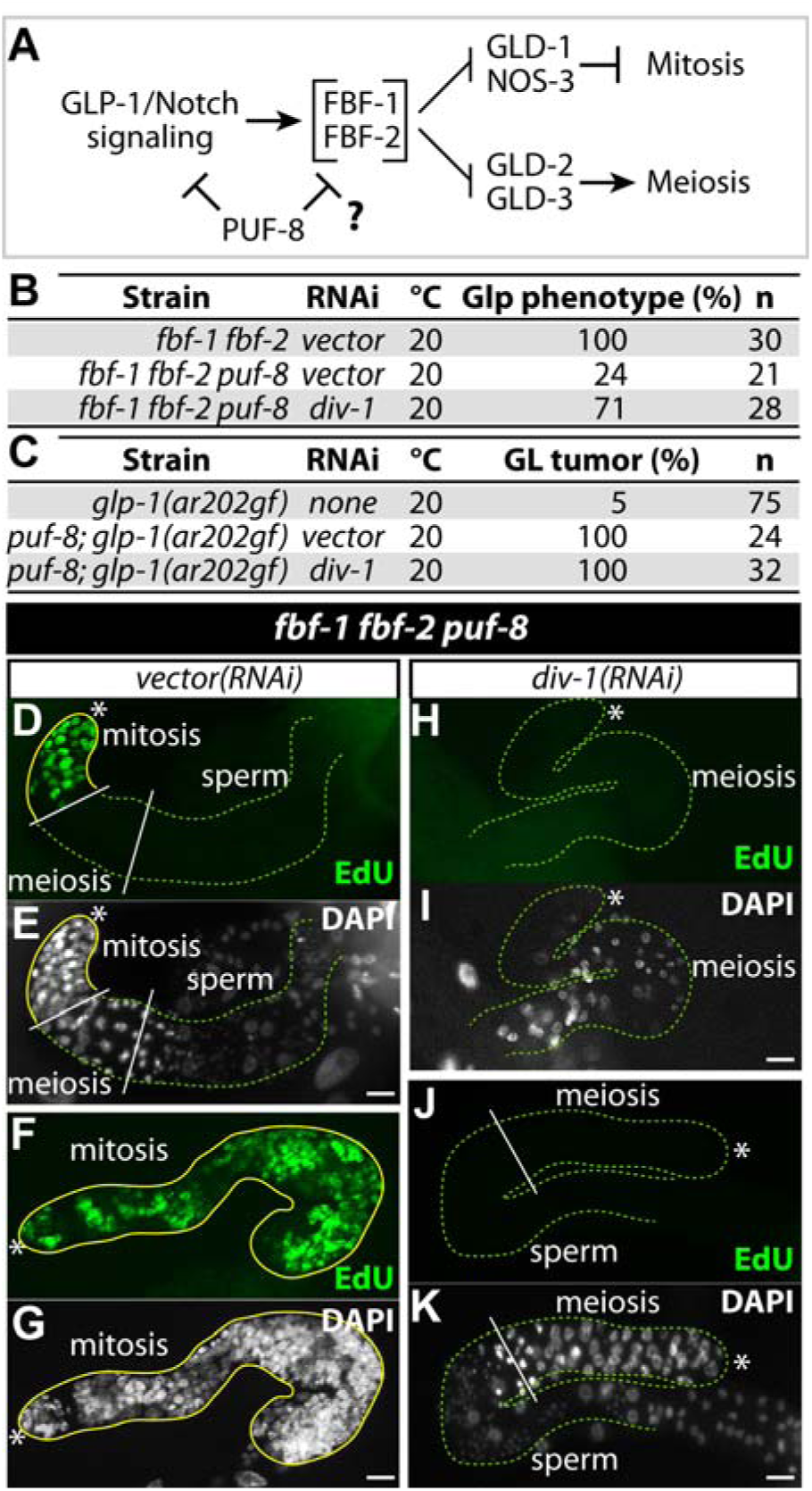
DIV-1 works together with FBF to promote GSC maintenance and germline proliferation. (A) A simplified network controlling the mitosis/meiosis decision. GLP-1/Notch signaling promotes GSC maintenance and germline proliferation, in part by transcriptional activation of the *fbf-2* gene (Lamont et al., 2004). FBF-1 and FBF-2 maintain GSCs by repressing the activity of the GLD-1/NOS-3 and GLD-2/GLD-3 pathways. Specifically, GLD-1/NOS-3 represses mitosis-promoting mRNAs and GLD-2/GLD-3 activates meiosis-promoting mRNAs. PUF-8 protein represses the proliferative fate through inhibiting GLP-1/Notch signaling (Racher and Hansen, 2012) and FBF (this study). (B) The percentage of the Glp phenotype. Glp phenotype was defined as germlines with no EdU-positive cells (see H-K). (C) The percentage of germline (GL) tumor phenotype. (D-G) EdU staining of dissected adult hermaphrodite germlines. *vector(RNAi); fbf-1 fbf-2 puf-8* mutants had mitotically dividing germ cells in the distal gonads (D and E) or throughout the germlines (F and G). (H-K) About 70% of *div-1(RNAi); fbf-1 fbf-2 puf-8* mutants lost mitotically dividing germ cells and displayed a Glp phenotype. EdU(-) germlines had two phenotypes; Glp germlines with only a few of meiotic germ cells (H and I) and with a few of meiotic germ cells and sperm (J and K). Solid yellow lines, a region with EdU(+) cells. Broken green lines, a region differentiating cells (meiotic cells and sperm). *, distal end. Scale bars: 10 µM.

It was previously reported that PUF-8 also represses GLP-1/Notch-mediated germline proliferation (Racher and Hansen, 2012; Datla et al., 2014) (Fig. 5A): while *glp-1(ar202gf)* mutants produce both sperm and oocyte, which are self-fertile at 20°C, the *puf-8* mutation strongly enhances the germline tumor phenotype of the *glp-1(ar202gf)* mutant even at 20°C (Racher and Hansen, 2012) (Fig. 5C). This suggests that PUF-8 inhibits proliferative fate through negative regulating GLP-1/Notch signaling or by functioning parallel to it (Racher and Hansen, 2012). We also tested if *div-1(RNAi)* suppresses the formation of germline tumors in *puf-8(q725); glp-1(ar202gf)* double mutants at 20°C. Interestingly, *div-1(RNAi)* failed to suppress the formation of germline tumors in *puf-8(q725); glp-1(ar202gf)* mutants (Fig. 5C). How does *div-1(RNAi)* suppress the formation of germline tumors of *glp-1(ar202gf)*, but not that of *puf-8; glp-1(ar202)*? One possible idea is that DIV-1 may fall the threshold for germline proliferation/differentiation decision, resulting in partially suppression of germline tumor phenotype (Fig. 7B, Supplementary material Fig. S5). However, *div-1(RNAi)* may be not sufficient to fall below the threshold that was raised by loss of PUF-8 (Supplementary material Fig. S5). These results suggest that DIV-1 and PUF-8 may have opposite roles in altering the threshold for GLP-1/Notch-mediated germline proliferation.

### DIV-1 is required for glp-1(q35) multi-vulva (Muv) formation

We showed here that DIV-1 promotes GLP-1/Notch-mediated germline proliferation. However, it still has a possibility that DIV-1 may influence germline proliferation through general cell cycle control, regardless of GLP-1/Notch signaling pathway. To test this possibility, we employed *glp-1(q35)* mutant. The *glp-1(q35)* has a premature stop codon, which results in a truncated GLP-1/Notch protein that lacks a negative regulatory domain (ProGluSerThr [PEST]) (Fig. 1D) (Mango et al., 1991). The PEST domain is associated with a shorter GLP-1(intra) half-life and more rapid degradation. The *glp-1(q35)* mRNA with a premature stop codon is degraded by nonsense-mediated decay (NMD) (Mango et al., 1991). Mutation of *smg-1*, which is required for NMD in *C. elegans*, stabilized *glp-1(q35)* mRNA, and thereby, suppressed the Glp phenotype of *glp-1(q35)* mutant. Wild-type adult hermaphrodites have normally one vulva (Fig. 6A and 6D). However, *smg-1; glp-1(q35)* double mutants have a multivulva phenotype similar to that caused by dominant *lin-12* (one of two *C. elegans* Notch receptors) mutations (Mango et al., 1991) (Fig. 1G, 6B and 6E). Next, to test whether DIV-1 is necessary for the *smg-1; glp-1(q35)* Muv phenotype, we performed RNAi of *div-1* and *vector* (negative control) into *smg-1; glp-1(q35)* double mutants at 20°C, and scored the number of extra vulva under differential interference contrast microscopy. Adult wild-type hermaphrodites typically have one vulva (Fig. 6A and 6D), but *vector(RNAi); smg-1; glp-1(q35)* mutants displayed a semidominant multivulva phenotype as well as the recessive, loss-of-function GLP phenotype (data not shown) (Mango et al., 1991). However, *div-1(RNAi)* significantly reduced the number of extra vulvae (Fig. 6C and 6F) and promote the Glp phenotype (data not shown) in the *smg-1; glp-1(q35)* mutants. We also examined the effects of other cell cycle regulators on *glp-1(q35)*-mediated muv phenotype. Intriguingly, depletion of CYE-1 and CYA-1 that showed an important role in GLP-1/Notch-mediated germline proliferation and germline tumor formation, respectively (Supplementary material Fig. S3) did not influence *glp-1(q35)*-mediated Muv formation (Supplementary material Fig. S4). By contrast, PRI-1 (a homolog of the DNA polymerase alpha-primase subunit B) appears to have a critical role in *glp-1(q35)*-mediated Muv formation (Supplementary material Fig. S4), although it did not play an important role in GLP-1/Notch-mediated germline proliferation and tumorigenesis, as previously reported (Fox et al., 2011) (Fig. 2E, Supplementary material Fig. S3). This result suggests that unique cell cycle length and structure may specify cell fate differently in germline and soma. All together, we propose that DIV-1 plays an important role in GLP-1/Notch signaling to control cellular events in the *C. elegans* germline and soma.

**Figure 6.**
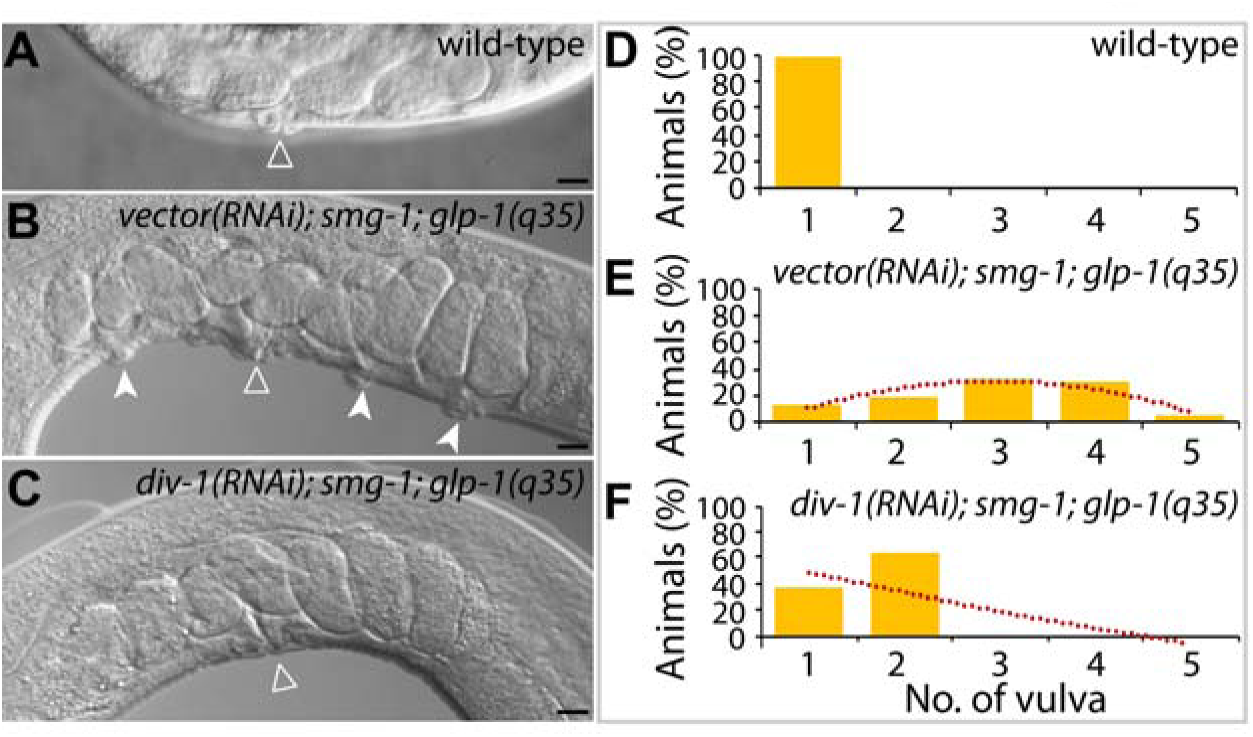
Depletion of DIV-1 suppresses the formation of multi-vulva caused by GLP-1(q35) activity. (A) Adult wild-type, Nomarski micrograph. A single vulva is induced (open triangle). (B) Adult *vector(RNAi); smg-1; glp-1(q35)* mutant, Nomarski micrograph. Ectopic vulvae (filled arrowhead) are induced in addition to the main vulva (open triangle). (C) Adult *div-1(RNAi); smg-1; glp-1(q35)* mutant, Nomarski micrograph. *div-1(RNAi)* suppress multi-vulva phenotype. (D-F) Graph showing the percentage of multi-vulva in adult animals. The number of vulva was determined by Nomarski DIC optics. Dotted red lines indicate trendline. See Supplementary material Fig. S4 for the effects of other cell cycle regulators on *glp-1(q35)*-mediated multi-vulva phenotype. Scale bars: 10 µM.

## DISCUSSION

In eukaryotes, intercellular signaling through Notch receptors regulates growth and differentiation during animal development (Lai, 2004). Moreover, aberrant regulation of the Notch signaling is highly associated with human diseases, including cancers (Allenspach et al., 2002; Rizzo et al., 2008; Yuan et al., 2015). In *C. elegans*, the mechanisms of GSC maintenance and germline proliferation largely rely on GLP-1/Notch signaling (Austin and Kimble, 1987) and a battery of RNA regulators (e.g., FBF (Kimble and Crittenden, 2007)). In this study, we found new tenable links between DIV-1 and GLP-1/Notch signaling in GSC maintenance, and germline proliferation. Importantly, stem cells have a unique cell cycle structure (short or absent G1 and long S phase), suggesting the potential role of G1/S and S phase in stem cell regulation. We specifically demonstrated for the first time that DIV-1 works together with GLP-1/Notch signaling and its downstream RNA regulators (e.g., FBF) for GSC maintenance and germline proliferation in *C. elegans*. All regulators that we studied here are highly conserved in other multicellular organisms, including humans. Therefore, our findings may provide insights into the cell cycle control of Notch-mediated cellular events in other model systems.

### Notch signaling and cell cycle control in stem cells and cancer cells

Notch signaling plays varied and critical roles in regulating many types of stem cells by receiving signals from their microenvironment (also known as “niche”). Notch signaling is critical in stem cells, not only for appropriately balancing self-renewal and differentiation, but also for regulating cell cycle progression to protect from tumorigenesis. In addition to Notch signaling, cell cycle control is a critical step in generating specific cells and tissues during development. The association of Notch signaling and cell cycle progression with cancer development has also been studied in various animal systems. For example, Notch signaling activates Cyclin D1 transcription and CDK2 activity, promoting S phase entry (Ronchini and Capobianco, 2001). In the pancreatic cancer cells, Notch signaling regulates the expression levels of Cyclin D1, Cyclin A, and cell cycle inhibitors (e.g., p21^CIP^ and p27^KIP1^, and CDK inhibitors) (Wang et al., 2006). In addition, Notch signaling mediates G1/S cell cycle progression through overexpression of Cyclin D3 and CDK4 in T cells (Joshi et al., 2009) and through reduction of p27^KIP1^, leading to more rapid cell cycle progression in T-cell acute lymphoblastic leukemia cells (Dohda et al., 2007). To date, many reports support the notion that Notch signaling affects cell cycle progression (Hori et al., 2013). However, it is poorly understood how cell cycle regulators work with Notch signaling to regulate cellular events in a whole animal. Recently, it was reported that Notch signaling is tightly linked to cell cycle progression during *C. elegans* vulval development (Nusser-Stein et al., 2012). Specifically, CYD-1/Cyclin D and CYE-1/Cyclin E stabilize LIN-12/Notch receptor in vulva precursor cells (VPCs) (Nusser-Stein et al., 2012). In *C. elegans* germline, CYE-1 is critical for GLP-1-mediated germline proliferation (Fox et al., 2011), but CYD-1 is not (Fox et al., 2011). Moreover, *C. elegans* CYD-1 and CYE-1 induce distinct cell cycle re-entry programs in differentiated muscle cells (Korzelius et al., 2011). These reports and our findings suggest that cell fate determination in different tissues (e.g., germline and soma) and/or developmental conditions (e.g., early/late development, early/terminal differentiation) may be regulated by different cell cycle progressions and quiescence. How does cell cycle regulate cell fate differently in germline and soma? We still do not know yet, but suggest that distinct cell cycle length and structure may contribute to cell fate specification differently in germline and soma. Similar features were observed in other systems, including mouse embryonic stem cells: greater than 70% of the cell population are in S phase (Savatier et al., 1996; Seidel and Kimble, 2015), suggesting that potential role of S phase in stem cell regulation.

### Role of DIV-1/PolA2 in GLP-1/Notch-mediated cellular events

The initiation of DNA replication during S phase relies on the DNA polymerase alpha-primase complex, including the catalytic subunit PolA1, the regulatory subunit PolA2, and the small and the large primase subunits “Prim1 and Prim2”, respectively. However, this complex seems to play additional roles in other cellular processes, such as DNA damage response, telomere maintenance, and the epigenetic control (Muzi-Falconi et al., 2003). This complex is well conserved in *C. elegans*: Y47D3A.29/PolA1, DIV-1/PolA2, PRI-1/Prim1, and PRI-2/Prim2. Our study showed that depletion of DIV-1/PolA2 affected GLP-1/Notch-mediated cellular events in the *C. elegans* germline and soma. How does DIV-1 control GLP-1/Notch-mediated GSC maintenance? We still do not know yet, but we suggest two mechanisms: *First*, DIV-1 may control GSC asymmetric division through cell division timing. A defining characteristic of stem cells is their ability to divide asymmetrically (Morrison and Kimble, 2006). Recently advanced studies provide ample evidence that many stem cells can still divide symmetrically, particularly when they are expanding in number during development or after injury (Morrison and Kimble, 2006). Thus, it is generally accepted that asymmetric division is a tool that stem cells can use to maintain stem cell homeostasis and appropriate numbers of progeny. Notably, *C. elegans div-1* was first identified as a key regulator for asymmetric cell division by controlling the proper timing of early embryonic cell division (Encalada et al., 2000). *div-1* mutants exhibit loss of asymmetry during early embryonic cleavages and result in the lacking of the endodermal and mesodermal cell fates (Encalada et al., 2000). This result speculates a possibility that *div-1* may also be involved in both asymmetric and symmetric division by controlling the proper timing of germ cell division. This idea can be determined by *in vivo* tracking of single cell behavior in the live *C. elegans* germline or by monitoring cell division in germ cell culture system. However, single germ cell tracking and germ cell culture systems have not yet been developed in *C. elegans*. Studying the role of *div-1* in the asymmetric/symmetric division of stem cells remains a challenge for the future. *Second*, depletion of *div-1* by RNAi may alter a threshold for germline proliferation/differentiation decision probably through cell division timing. As described in Fig. 7B, DIV-1/FBF and GLDs/NOS-3 may control the threshold for germline proliferation/differentiation decision. In addition, PUF-8/GLDs/NOS-3 may inhibit GLP-1 activity. Notch signaling controls cell division timing (Hunter et al., 2016). Therefore, Notch signaling, downstream regulator (PUF proteins), and DIV-1 may work together to control the threshold for germline proliferation/differentiation decision probably through cell division timing. This is consistent with the known role of DIV-1 in embryonic cell division timing (Encalada et al., 2000). However, understanding the mechanism of how aberrant cell division timing controls the threshold for germline proliferation/differentiation decision remains a major challenge for the future.

**Figure 7.**
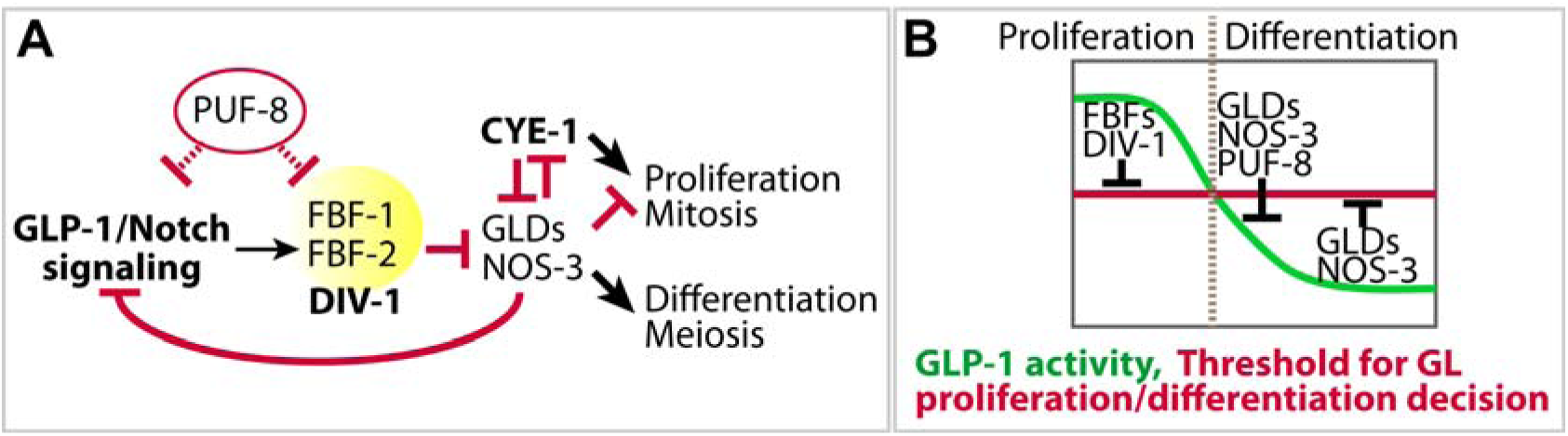
Working model. (A) DIV-1 works together with FBFs to promote germline proliferation and inhibit differentiation. PUF-8 inhibits the function of GLP-1/Notch signaling and its downstream regulators (FBFs and probably DIV-1). (B) The potential roles of DIV-1 and other key regulators in controlling a threshold for germline proliferation/differentiation decision. See Supplementary material Fig. S5 for the effects of other key regulators on controlling a threshold for germline proliferation/differentiation decision.

### DIV-1/PolA2 as a potential target for cancer treatments

PolA2 (a regulatory subunit of the DNA polymerase) plays an essential role in the initiation of DNA replication. During the S phase of the cell cycle, the PolA2 is recruited to DNA at the replicative forks. PolA2 (G583R) mutation led to localize in the cytoplasm instead of the nucleus, which inhibits DNA replication in cancer cells (e.g., non-small cell lung cancer (NSCLC) and makes them sensitive to chemotherapy (e.g., Gemcitabine) (Mah et al., 2014). In addition, the synthetic retinoid 6-[3-(1-adamantyl)-4-hydroxyphenyl]-2-naphthalene carboxylic acid (CD437) is an inhibitor of PolA1 and selectively induces apoptosis in human lung cancer cells (Sun et al., 2002). Cancer cells have a different response to inhibition of PolA1 by CD437 as compared to aphidicolin. CD437-mediated inhibition of PolA1 leads to cell death in cancer cells, but it induces cell cycle arrest in normal epithelial cells (Sun et al., 2002). Notably, Notch signaling is altered in approximately one third of NSCLCs, which are the leading cause of cancer-related deaths. Although the role of PolA1/2 in elevated Notch activity-mediated NSCLCs has not yet been studied, a fundamental mechanism in stem cell maintenance and tumorigenesis may be conserved. Therefore, understanding a mechanism underlying Notch signaling and PolA1/2 may hold promise for molecular and cellular therapies in these tumors.

## MATERIALS AND METHODS

### Nematode strains

All strains were derived from Bristol strain N2 and maintained at 20°C as described unless otherwise noticed (Brenner, 1974). Mutants and transgenic worms are listed in Supplementary Table S1.

### Hydroxyurea (HU) treatment and whole worm DAPI staining

L2/3 stage of wild-type and *glp-1(bn18)* worms were transferred to NGM (Nematode Growth Media) agar plates containing 40 mM Hydroxyurea (HU) and incubated at 20°C. After 18 hours with HU plate incubation, worms were transferred to a normal NGM plate and let them grow up to adults. For DAPI staining, whole worms were collected using M9 buffer from NGM agar plates and transferred into a 2 ml Eppendorf tube. After washing two times with M9 buffer (3 g KH_2_PO_4_, 6 g Na_2_HPO_4_, 5 g NaCl, 1 ml 1 M MgSO_4_, H_2_O to 1 litre), whole worms were fixed with 3% paraformaldehyde/0.1M K_2_HPO_4_ (pH 7.2) solution for 20 min, and then post-fixed with cold 100% methanol for 5 min at −20°C (alternatively, fixed worms can be stored in cold methanol at -20°C for a few days). After washing with 1x PTW (1x PBS and 0.1% Tween 20), add DAPI (100 ng/mL) and incubate the fixed gonads for 10 min at room temperature, following by washing them with 1x PTW three times. DAPI staining was observed using a fluorescence microscopy, as previously described (Yoon et al., 2016).

### RNA interference (RNAi)

RNAi constructs were obtained from the *C. elegans* RNAi Library (Thermo Scientific) or kindly provided by Kimble’s lab (University of Wisconsin-Madison). RNAi was performed by feeding adult staged worms (P0) bacterial expressing double stranded RNAs (dsRNAs) corresponding to the gene of interest as previously described (Kamath et al., 2001; Ashrafi et al., 2003) (see Fig. 2E). The germline phenotypes were analyzed when F1 progeny reach to adult stage (3 days later from L1). The effect of *div-1* and *cye-1* (positive control) on GSC maintenance and germline proliferation in adult *glp-1(bn18)* mutants was tested by RNAi from young adult animals (see Fig. 2J). 3 days (72 h) later, the germline phenotypes were analyzed by staining dissected gonads with anti-REC-8 and anti-HIM-3 antibodies.

### Germline antibody staining

Germline antibody staining was performed as described in (Yoon et al., 2016). Briefly, dissected gonads were fixed in 3% paraformaldehyde/0.1M K_2_HPO_4_ (pH 7.2) solution for 10-20 min, and then post-fixed with cold 100% methanol for 5 min (Alternatively, fixed worms can be stored in cold methanol at -20°C for a few days). After 30 min blocking with 1x PTW/0.5% BSA (Bovine Serum Albumin) solution, add primary antibody and incubate for 2 hours at room temperature or overnight at 4°C. The dissected gonads were washed three times for at least 30 min (10-min interval) with 1x PTW/0.5%BSA solution and incubated the dissected gonads in the 1x PTW/0.5% BSA solution containing the fluorescence-conjugated secondary antibodies for 1-2 hours at room temperature. After washing three times using 1x PTW/0.5% BSA solution for at least 30 min (10-min interval), the dissected gonads were stained with DAPI solution (100 ng/mL) for 10 min at room temperature and were then washed with 1x PTW/0.5% BSA solution three times. The antibody staining was observed using a fluorescence microscopy. See Supplementary Table 2 for a list and working conditions of primary antibodies that we used in this study.

### EdU labeling

For EdU labeling, animals were incubated with rocking in M9/0.1% Tween 20/1 mM EdU for 30 min at room temperature. Gonads were dissected as for germline antibody staining and fixed in 3% paraformaldehyde/0.1M K_2_HPO_4_ (pH 7.2) solution for 10-20 min, followed by -20°C methanol fixation for 10 min. Dissected gonads were blocked in 1xPTW/0.5% BSA solution for 30 min at room temperature. EdU labeling was detected using the Click-iT EdU Alexa Fluor 488 Imaging Kit (Invitrogen, CA, #C10337), according to the manufacturer’s instructions. After washing three times with 1x PTW/0.5% BSA solution for at least 30 min (10-min interval), the dissected gonads were stained with DAPI solution (100 ng/mL) for 10 min at room temperature and were washed with 1x PTW/0.5% BSA solution three times. The EdU labeling was observed using a fluorescence microscopy.

### S-phase index

S-phase index was determined by pulsing animals with EdU for 30 min and counting total cells of REC-8-positve cells, as previously described (Crittenden et al., 2006; Fox et al., 2011).

## ACKNOWLEDGEMENTS

We are grateful to Dr. Eleanor Maine, Dr. Brett D. Keiper, and Dr. Elizabeth T. Ables for advice and discussion during this work. We thank Dr. Judith Kimble (University of Wisconsin-Madison, HHMI) for *C. elegans* mutant strains and antibodies (anti-GLP-1 and anti-GLD-1) and Dr. Josef Loidl (University of Vienna) for REC-8 antibody. This work was supported in part by the Vidant Medical Center-Cancer Research and Education Fund, Brody Brother’s Grant (BBE 213152), Oncology Grant, BSOM Seed/Bridge Grant, and NIH (1R15GM112174-01A1) to M-H.L. The Caenorhabditis Genetics Center (CGC) is supported by the National Institutes of Health - Office of Research Infrastructure Programs (P40 OD010440).

## AUTHOR CONTRIBUTIONS

DSY, DSC, and MHL conceived and designed the experiments, analyzed the data, and wrote the paper. DSY, DSC, and MHL performed the experiments.

## REFERENCES

1. Ables, E. T. and Drummond-Barbosa, D. (2013) ‘Cyclin E controls Drosophila female germline stem cell maintenance independently of its role in proliferation by modulating responsiveness to niche signals’, Development 140(3): 530–40.

2. Allenspach, E. J., Maillard, I., Aster, J. C. and Pear, W. S. (2002) ‘Notch signaling in cancer’, Cancer Biol Ther 1(5): 466–76.

3. Ariz, M., Mainpal, R. and Subramaniam, K. (2009) ‘C. elegans RNA-binding proteins PUF-8 and MEX-3 function redundantly to promote germline stem cell mitosis’, Dev Biol 326(2): 295–304.

4. Ashrafi, K., Chang, F. Y., Watts, J. L., Fraser, A. G., Kamath, R. S., Ahringer, J. and Ruvkun, G. (2003) ‘Genome-wide RNAi analysis of Caenorhabditis elegans fat regulatory genes’, Nature 421(6920): 268–72.

5. Austin, J. and Kimble, J. (1987) ‘glp-1 is required in the germ line for regulation of the decision between mitosis and meiosis in C. elegans’, Cell 51(4): 589–99.

6. Berry, L. W., Westlund, B. and Schedl, T. (1997) ‘Germ-line tumor formation caused by activation of glp-1, a Caenorhabditis elegans member of the Notch family of receptors’, Development 124(4): 925–36.

7. Besson, A., Dowdy, S. F. and Roberts, J. M. (2008) ‘CDK inhibitors: cell cycle regulators and beyond’, Dev Cell 14(2): 159–69.

8. Brenner, S. (1974) ‘The genetics of Caenorhabditis elegans’, Genetics 77(1): 71–94.

9. Cha, D. S., Datla, U. S., Hollis, S. E., Kimble, J. and Lee, M. H. (2012) ‘The Ras-ERK MAPK regulatory network controls dedifferentiation in Caenorhabditis elegans germline’, Biochim Biophys Acta 1823(10): 1847–55.

10. Crittenden, S. L., Bernstein, D. S., Bachorik, J. L., Thompson, B. E., Gallegos, M., Petcherski, A. G., Moulder, G., Barstead, R., Wickens, M. and Kimble, J. (2002) ‘A conserved RNA-binding protein controls germline stem cells in Caenorhabditis elegans’, Nature 417(6889): 660–3.

11. Crittenden, S. L., Leonhard, K. A., Byrd, D. T. and Kimble, J. (2006) ‘Cellular analyses of the mitotic region in the Caenorhabditis elegans adult germ line’, Mol Biol Cell 17(7): 3051–61.

12. Datla, U. S., Scovill, N. C., Brokamp, A. J., Kim, E., Asch, A. S. and Lee, M. H. (2014) ‘Role of PUF-8/PUF protein in stem cell control, sperm-oocyte decision and cell fate reprogramming’, J Cell Physiol 229(10): 1306–11.

13. Dohda, T., Maljukova, A., Liu, L., Heyman, M., Grander, D., Brodin, D., Sangfelt, O. and Lendahl, U. (2007) ‘Notch signaling induces SKP2 expression and promotes reduction of p27Kip1 in T-cell acute lymphoblastic leukemia cell lines’, Exp Cell Res 313(14): 3141–52.

14. Eckmann, C. R., Crittenden, S. L., Suh, N. and Kimble, J. (2004) ‘GLD-3 and control of the mitosis/meiosis decision in the germline of Caenorhabditis elegans’, Genetics 168(1): 147–60.

15. Encalada, S. E., Martin, P. R., Phillips, J. B., Lyczak, R., Hamill, D. R., Swan, K. A. and Bowerman, B. (2000) ‘DNA replication defects delay cell division and disrupt cell polarity in early Caenorhabditis elegans embryos’, Dev Biol 228(2): 225–38.

16. Fox, P. M. and Schedl, T. (2015) ‘Analysis of Germline Stem Cell Differentiation Following Loss of GLP-1 Notch Activity in Caenorhabditis elegans’, Genetics 201(1): 167–84.

17. Fox, P. M., Vought, V. E., Hanazawa, M., Lee, M. H., Maine, E. M. and Schedl, T. (2011) ‘Cyclin E and CDK-2 regulate proliferative cell fate and cell cycle progression in the C. elegans germline’, Development 138(11): 2223–34.

18. Girard, F., Strausfeld, U., Fernandez, A. and Lamb, N. J. (1991) ‘Cyclin A is required for the onset of DNA replication in mammalian fibroblasts’, Cell 67(6): 1169–79.

19. Hansen, D., Hubbard, E. J. and Schedl, T. (2004a) ‘Multi-pathway control of the proliferation versus meiotic development decision in the Caenorhabditis elegans germline’, Dev Biol 268(2): 342–57.

20. Hansen, D., Wilson-Berry, L., Dang, T. and Schedl, T. (2004b) ‘Control of the proliferation versus meiotic development decision in the C. elegans germline through regulation of GLD-1 protein accumulation’, Development 131(1): 93–104.

21. Henderson, S. T., Gao, D., Lambie, E. J. and Kimble, J. (1994) ‘lag-2 may encode a signaling ligand for the GLP-1 and LIN-12 receptors of C. elegans’, Development 120(10): 2913–24.

22. Hori, K., Sen, A. and Artavanis-Tsakonas, S. (2013) ‘Notch signaling at a glance’, J Cell Sci 126(Pt 10): 2135–40.

23. Hubbard, E. J. (2007) ‘Caenorhabditis elegans germ line: a model for stem cell biology’, Dev Dyn 236(12): 3343–57.

24. Hunter, G. L., Hadjivasiliou, Z., Bonin, H., He, L., Perrimon, N., Charras, G. and Baum, B. (2016) ‘Coordinated control of Notch/Delta signalling and cell cycle progression drives lateral inhibition-mediated tissue patterning’, Development 143(13): 2305–10.

25. Johnson, D. G. and Walker, C. L. (1999) ‘Cyclins and cell cycle checkpoints’, Annu Rev Pharmacol Toxicol 39: 295–312.

26. Joshi, I., Minter, L. M., Telfer, J., Demarest, R. M., Capobianco, A. J., Aster, J. C., Sicinski, P., Fauq, A., Golde, T. E. and Osborne, B. A. (2009) ‘Notch signaling mediates G1/S cell-cycle progression in T cells via cyclin D3 and its dependent kinases’, Blood 113(8): 1689–98.

27. Kadyk, L. C. and Kimble, J. (1998) ‘Genetic regulation of entry into meiosis in Caenorhabditis elegans’, Development 125(10): 1803–13.

28. Kalchhauser, I., Farley, B. M., Pauli, S., Ryder, S. P. and Ciosk, R. (2011) ‘FBF represses the Cip/Kip cell-cycle inhibitor CKI-2 to promote self-renewal of germline stem cells in C. elegans’, EMBO J 30(18): 3823–9.

29. Kamath, R. S., Martinez-Campos, M., Zipperlen, P., Fraser, A. G. and Ahringer, J. (2001) ‘Effectiveness of specific RNA-mediated interference through ingested double-stranded RNA in Caenorhabditis elegans’, Genome Biol 2(1): RESEARCH0002.

30. Kennedy, S., Wang, D. and Ruvkun, G. (2004) ‘A conserved siRNA-degrading RNase negatively regulates RNA interference in C. elegans’, Nature 427(6975): 645–9.

31. Kershner, A. M., Shin, H., Hansen, T. J. and Kimble, J. (2014) ‘Discovery of two GLP-1/Notch target genes that account for the role of GLP-1/Notch signaling in stem cell maintenance’, Proc Natl Acad Sci U S A 111(10): 3739–44.

32. Kimble, J. and Crittenden, S. L. (2007) ‘Controls of germline stem cells, entry into meiosis, and the sperm/oocyte decision in Caenorhabditis elegans’, Annu Rev Cell Dev Biol 23: 405–33.

33. Kobet, R. A., Pan, X., Zhang, B., Pak, S. C., Asch, A. S. and Lee, M. H. (2014) ‘Caenorhabditis elegans: A Model System for Anti-Cancer Drug Discovery and Therapeutic Target Identification’, Biomol Ther (Seoul) 22(5): 371–83.

34. Kodoyianni, V., Maine, E. M. and Kimble, J. (1992) ‘Molecular basis of loss-of-function mutations in the glp-1 gene of Caenorhabditis elegans’, Mol Biol Cell 3(11): 1199–213.

35. Korzelius, J., The, I., Ruijtenberg, S., Prinsen, M. B., Portegijs, V., Middelkoop, T. C., Groot Koerkamp, M. J., Holstege, F. C., Boxem, M. and van den Heuvel, S. (2011) ‘Caenorhabditis elegans cyclin D/CDK4 and cyclin E/CDK2 induce distinct cell cycle re-entry programs in differentiated muscle cells’, PLoS Genet 7(11): e1002362.

36. Lai, E. C. (2004) ‘Notch signaling: control of cell communication and cell fate’, Development 131(5): 965–73.

37. Lamont, L. B., Crittenden, S. L., Bernstein, D., Wickens, M. and Kimble, J. (2004) ‘FBF-1 and FBF-2 regulate the size of the mitotic region in the C. elegans germline’, Dev Cell 7(5): 697–707.

38. Lee, M. H., Hook, B., Lamont, L. B., Wickens, M. and Kimble, J. (2006) ‘LIP-1 phosphatase controls the extent of germline proliferation in Caenorhabditis elegans’, EMBO J 25(1): 88–96.

39. Liu, J., Sato, C., Cerletti, M. and Wagers, A. (2010) ‘Notch signaling in the regulation of stem cell self-renewal and differentiation’, Curr Top Dev Biol 92: 367–409.

40. Mah, T. L., Yap, X. N., Limviphuvadh, V., Li, N., Sridharan, S., Kuralmani, V., Feng, M., Liem, N., Adhikari, S., Yong, W. P. et al. (2014) ‘Novel SNP improves differential survivability and mortality in non-small cell lung cancer patients’, BMC Genomics 15 Suppl 9: S20.

41. Maine, E. M., Hansen, D., Springer, D. and Vought, V. E. (2004) ‘Caenorhabditis elegans atx-2 promotes germline proliferation and the oocyte fate’, Genetics 168(2): 817–30.

42. Mango, S. E., Maine, E. M. and Kimble, J. (1991) ‘Carboxy-terminal truncation activates glp-1 protein to specify vulval fates in Caenorhabditis elegans’, Nature 352(6338): 811–5.

43. Morrison, S. J. and Kimble, J. (2006) ‘Asymmetric and symmetric stem-cell divisions in development and cancer’, Nature 441(7097): 1068–74.

44. Murray, A. W. (2004) ‘Recycling the cell cycle: cyclins revisited’, Cell 116(2): 221–34.

45. Muzi-Falconi, M., Giannattasio, M., Foiani, M. and Plevani, P. (2003) ‘The DNA polymerase alpha-primase complex: multiple functions and interactions’, ScientificWorldJournal 3: 21–33.

46. Nusser-Stein, S., Beyer, A., Rimann, I., Adamczyk, M., Piterman, N., Hajnal, A. and Fisher, J. (2012) ‘Cell-cycle regulation of NOTCH signaling during C. elegans vulval development’, Mol Syst Biol 8: 618.

47. Ohtsubo, M., Theodoras, A. M., Schumacher, J., Roberts, J. M. and Pagano, M. (1995) ‘Human cyclin E, a nuclear protein essential for the G1-to-S phase transition’, Mol Cell Biol 15(5): 2612–24.

48. Pepper, A. S., Killian, D. J. and Hubbard, E. J. (2003) ‘Genetic analysis of Caenorhabditis elegans glp-1 mutants suggests receptor interaction or competition’, Genetics 163(1): 115–32.

49. Priti, A. and Subramaniam, K. (2015) ‘PUF-8 Functions Redundantly with GLD-1 to Promote the Meiotic Progression of Spermatocytes in Caenorhabditis elegans’, G3 (Bethesda) 5(8): 1675–84.

50. Pushpa, K., Kumar, G. A. and Subramaniam, K. (2013) ‘PUF-8 and TCER-1 are essential for normal levels of multiple mRNAs in the C. elegans germline’, Development 140(6): 1312–20.

51. Qiao, L., Lissemore, J. L., Shu, P., Smardon, A., Gelber, M. B. and Maine, E. M. (1995) ‘Enhancers of glp-1, a gene required for cell-signaling in Caenorhabditis elegans, define a set of genes required for germline development’, Genetics 141(2): 551–69.

52. Racher, H. and Hansen, D. (2012) ‘PUF-8, a Pumilio homolog, inhibits the proliferative fate in the Caenorhabditis elegans germline’, G3 (Bethesda) 2(10): 1197–205.

53. Rizzo, P., Osipo, C., Foreman, K., Golde, T., Osborne, B. and Miele, L. (2008) ‘Rational targeting of Notch signaling in cancer’, Oncogene 27(38): 5124–31.

54. Ronchini, C. and Capobianco, A. J. (2001) ‘Induction of cyclin D1 transcription and CDK2 activity by Notch(ic): implication for cell cycle disruption in transformation by Notch(ic)’, Mol Cell Biol 21(17): 5925–34.

55. Savatier, P., Lapillonne, H., van Grunsven, L. A., Rudkin, B. B. and Samarut, J. (1996) ‘Withdrawal of differentiation inhibitory activity/leukemia inhibitory factor up-regulates D-type cyclins and cyclin-dependent kinase inhibitors in mouse embryonic stem cells’, Oncogene 12(2): 309–22.

56. Seidel, H. S. and Kimble, J. (2015) ‘Cell-cycle quiescence maintains Caenorhabditis elegans germline stem cells independent of GLP-1/Notch’, Elife 4.

57. Sorokin, E. P., Gasch, A. P. and Kimble, J. (2014) ‘Competence for chemical reprogramming of sexual fate correlates with an intersexual molecular signature in Caenorhabditis elegans’, Genetics 198(2): 561–75.

58. Subramaniam, K. and Seydoux, G. (2003) ‘Dedifferentiation of primary spermatocytes into germ cell tumors in C. elegans lacking the pumilio-like protein PUF-8’, Curr Biol 13(2): 134–9.

59. Sun, S. Y., Yue, P., Chen, X., Hong, W. K. and Lotan, R. (2002) ‘The synthetic retinoid CD437 selectively induces apoptosis in human lung cancer cells while sparing normal human lung epithelial cells’, Cancer Res 62(8): 2430–6.

60. Vaid, S., Ariz, M., Chaturbedi, A., Kumar, G. A. and Subramaniam, K. (2013) ‘PUF-8 negatively regulates RAS/MAPK signalling to promote differentiation of C. elegans germ cells’, Development 140(8): 1645–54.

61. Vermeulen, K., Van Bockstaele, D. R. and Berneman, Z. N. (2003) ‘The cell cycle: a review of regulation, deregulation and therapeutic targets in cancer’, Cell Prolif 36(3): 131–49.

62. Wang, Z., Zhang, Y., Li, Y., Banerjee, S., Liao, J. and Sarkar, F. H. (2006) ‘Down-regulation of Notch-1 contributes to cell growth inhibition and apoptosis in pancreatic cancer cells’, Mol Cancer Ther 5(3): 483–93.

63. Wickens, M., Bernstein, D. S., Kimble, J. and Parker, R. (2002) ‘A PUF family portrait: 3’UTR regulation as a way of life’, Trends Genet 18(3): 150–7.

64. Wong, M. D., Jin, Z. and Xie, T. (2005) ‘Molecular mechanisms of germline stem cell regulation’, Annu Rev Genet 39: 173–95.

65. Yoon, D. S., Pendergrass, D. L. and Lee, M. H. (2016) ‘A simple and rapid method for combining fluorescent in situ RNA hybridization (FISH) and immunofluorescence in the C. elegans germline’, MethodsX 3: 378–85.

66. Yuan, X., Wu, H., Xu, H., Xiong, H., Chu, Q., Yu, S., Wu, G. S. and Wu, K. (2015) ‘Notch signaling: an emerging therapeutic target for cancer treatment’, Cancer Lett 369(1): 20–7.

67. Zetka, M. C., Kawasaki, I., Strome, S. and Muller, F. (1999) ‘Synapsis and chiasma formation in Caenorhabditis elegans require HIM-3, a meiotic chromosome core component that functions in chromosome segregation’, Genes Dev 13(17): 2258–70.

